# Pro-apoptotic caspase deficiency reveals a cell-extrinsic mechanism of NK cell regulation

**DOI:** 10.1101/2021.09.09.459676

**Authors:** Tayla M. Olsen, Wei Hong Tan, Arne C. Knudsen, Anthony Rongvaux

## Abstract

Regulated cell death is essential for the maintenance of cellular and tissue homeostasis. In the hematopoietic system, genetic defects in apoptotic cell death generally produce the accumulation of immune cells, inflammation and autoimmunity. In contrast, we found that genetic deletion of caspases of the mitochondrial apoptosis pathway reduces natural killer (NK) cell numbers and makes NK cells functionally defective *in vivo* and *in vitro*. Caspase deficiency results in constitutive activation of a type I interferon (IFN) response, due to leakage of mitochondrial DNA and activation of the cGAS/STING pathway. The NK cell defect in caspase-deficient mice is independent of the type I IFN response, but the phenotype is partially rescued by cGAS or STING deficiency. Finally, caspase deficiency alters NK cells in a cell-extrinsic manner. Type I IFNs and NK cells are two essential effectors of antiviral immunity, and our results demonstrate that they are both regulated in a caspase-dependent manner. Beyond caspase-deficient animals, our observations may have implications in infections that trigger mitochondrial stress and caspase-dependent cell death.

## Introduction

Cellular homeostasis in the immune system is maintained by a balance between the development of new cells from hematopoietic progenitors [1], and the elimination of surplus cells by apoptotic cell death [2, 3]. The mitochondrial pathway of apoptosis plays a central role in immune cell death [2]. It is controlled by Bcl-2 family members, which regulate Bax/Bak pore formation and mitochondrial outer membrane permeabilization (MOMP). Transgenic overexpression of the anti-apoptotic Bcl-2 protein, or deficiency in pro-apoptotic Bcl-2 family members, such as Bim, result in defective apoptotic cell death, immune cell hyperplasia, and autoimmune disorders [4-6]. MOMP is generally considered a point of no return in the initiation of mitochondrial apoptosis [7, 8].

MOMP enables the release of cytochrome *c* from the mitochondrial intermembrane space to the cytosol, where it assembles with Apaf-1 and the initiator caspase-9, in a multiprotein complex called the apoptosome [9]. Activation of caspase-9 in this complex triggers cleavage and activation of downstream effectors, caspase-3 and caspase-7, resulting in apoptotic demise of the cell [10]. However, there are exceptions to this canonical Bcl-2/apoptosome/caspases model of mitochondrial cell death. Indeed, in most immune cells, mitochondrial cell death is triggered independently of Apaf-1 and caspase-9 – unlike Bcl-2 transgenic or Bim-deficient mice, mice lacking Apaf-1 or caspase-9 display almost normal lymphocyte cellularity [11].

Although pro-apoptotic caspases are dispensable for mitochondria-initiated cell death [11], they are conserved across all metazoans [12]. This raises the question of which selective advantages are conferred by caspases, resulting in their evolutionary conservation as cell death mediators. It has been postulated that pro-apoptotic caspases uniquely kill cells in an “immunologically silent” manner [13, 14], but the mechanisms by which they silence inflammatory responses are diverse and incompletely described. These mechanisms include the inactivation of intracellular proteins, known as damage-associated molecular patterns (DAMPs). that would otherwise trigger inflammation upon release into the extracellular milieu, such as HMGB1 or interleukin (IL)-33 [15, 16]. Another anti-inflammatory mechanism of pro-apoptotic caspases is the induction of immune tolerance toward cytosolic mitochondrial DAMPs [17]. Mitochondria originated from an endosymbiotic event and retain molecular patterns from their bacterial ancestry, capable of engaging innate immune sensors [18, 19]. Upon Bax/Bak-dependent MOMP, mitochondrial DNA (mtDNA) is exposed to the nucleic acid sensor, cGAS, and triggers cell-intrinsic inflammation through the STING pathway and type I interferon (IFN) expression [20-23]. However, this process is negatively regulated by caspases, ensuring the immunologically silent nature of caspase-mediated apoptosis [20, 21].

Mice lacking pro-apoptotic caspases uncover immunoregulatory mechanisms that are controlled by caspases in normal conditions. We postulate that these mechanisms can be activated in pathologies or stress conditions that bypass caspase activation. For example, caspase-deficient mice were previously used to reveal the existence of the mtDNA/cGAS/STING axis, based on a broad state of IFN-mediated viral resistance induced in vivo by caspase deficiency [20, 21]. A role for mtDNA-dependent activation of cGAS/STING was subsequently recognized in several pathological conditions, including viral and bacterial infection [24-28], as well as inflammatory diseases [29, 30].

Natural Killer (NK) cells are innate lymphoid cells specialized in the elimination of abnormal cells, such as infected and/or malignant cells [31]. NK cell effector functions include the direct killing of target cells, as well as the production of cytokines, such as interferon (IFN)-γ [32]. The relevance of NK cells in human diseases is demonstrated by epidemiological and genetic studies, which have revealed associations between NK cell deficiencies and increased incidence of various types of cancer or susceptibility to chronic viral infections [33, 34]. Here, we identify a defect in NK cell development and function in mice lacking pro-apoptotic caspases. Our work reveals a novel regulatory mechanism of NK cell development and function, which may be relevant in conditions of cellular stress or cell death that bypass pro-apoptotic caspases.

## Results

### NK cell numbers are reduced in caspase-deficient mice

Conditional deletion, with the Tie2-Cre(E+H) transgene [35], of caspase-9 (*Casp9*^*fl/fl*^ *Tie2-Cre*^*+*^) or of caspase-3 in caspase-7-deficient background (*Casp3*^*fl/fl*^ *Casp7*^*-/-*^ *Tie2-Cre*^*+*^) results in mice lacking effectors of the mitochondrial apoptosis pathway in immune cells [21]. Unlike Bcl-2 overexpression (**Supplementary Figure 1A**) [4, 5] or Bim-deficiency [6], caspase-deficiency doesn’t significantly affect immune cell numbers (**Supplementary Figure 1A**). This observation is in agreement with a previous report [11], and confirms that caspases are dispensable for Bcl2-regulated cell death and immune cell homeostasis. The overall lineage composition of immune cells is only mildly affected by caspase deficiency (**Supplementary Figure 1B**). However, we noticed a 2.5- to 4-fold reduction in the frequency and absolute numbers of NK cells in the spleen and blood of *Casp9*^*fl/fl*^ *Tie2-Cre*^*+*^ and *Casp3*^*fl/fl*^ *Casp7*^*-/-*^ *Tie2-Cre*^*+*^ mice, compared to their *Tie2-Cre*^*-*^ littermates (**Supplementary Figure 1C, Figure 1A-C**). We also compared homozygous *Casp9*^*fl/fl*^ *Tie2-Cre*^*+*^ mice to their heterozygous *Casp9*^*+/fl*^ *Tie2-Cre*^*+*^ littermates (**Supplementary Figure 1D**), and we generated hematopoietic chimeras by transplantation of fetal liver cells from *Casp9*^*+/+*^ and *Casp9*^*Δ/Δ*^ littermate embryos into irradiated wildtype congenic recipient mice (**Supplementary Figure 1E**). Both experiments confirmed the NK cell defect in the absence of caspase-9. Together, these results demonstrate the hematopoietic origin of the phenotype, and exclude a potential artefactual role of the *Tie2-Cre* transgene.

**Figure 1.**
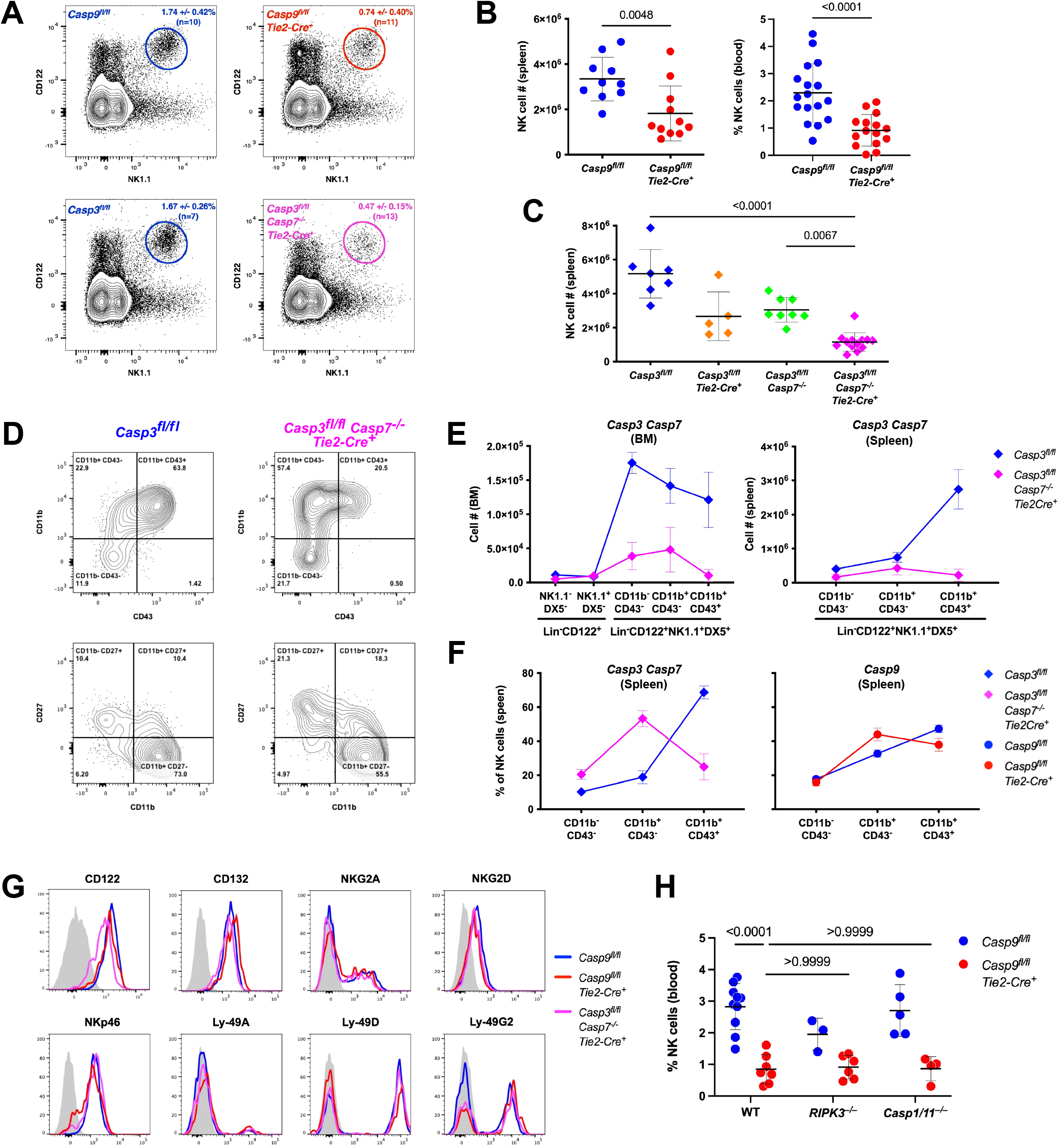
Absence of caspases of the intrinsic apoptosis pathway results in reduced NK cell numbers. (**A**-**C**) Mice with conditional deletion, in their hematopoietic system, of the initiator caspase-9 (*Casp9*^fl/fl^ *Tie2-Cre*^+^) or of the downstream effector caspases (*Casp3*^fl/fl^ *Casp7*^-/-^ *Tie2-Cre*^+^) have reduced frequencies and numbers of NK cells in the blood and spleen. (**A**) Representative flow cytometry analysis of splenic NK cells, identified as CD122^+^ NK1.1^+^ cells. Numbers indicate mean +/- S.D. of cell percentage in the gate. (**B**) Absolute numbers in the spleen and frequency in the blood of NK cells identified as Lin^-^ CD122^+^ NK1.1^+^ DX5^+^ cells in mice lacking caspase-9 (N=10-17 mice per genotype). (**C**) Absolute numbers of NK cells in the spleen of mice lacking caspase-3 and caspase-7 (N=5-13 mice per genotype). Results are combined from at least two independent experiments, including male and female mice. Error bars indicate mean +/- S.D. P-values calculated with Mann-Whitney or Kruskal-Wallis tests. (**D**-**F**) NK cells in caspase-deficient mice have an immature phenotype, as determined by the expression of CD11b, CD43 and CD27. (**D**) Representative flow cytometry plots from splenic NK cells (Lin^-^ CD122^+^ NK1.1^+^ DX5^+^). (**E**) Absolute numbers of NK cells at distinct maturation stages in the bone marrow and spleen in the spleen. (**F**) Frequency of NK cell maturation stages in the spleen. Error bars indicate mean +/- S.D. of 3-5 mice, representative of at least two independent experiments. (**G**) Expression of cell surface markers by NK cells, identified as Lin^-^ NK1.1^+^ DX5^+^ cells. The grey histogram represents FMO staining of the wildtype (*Casp9*^*fl/fl*^) sample. (**H**) Frequency of NK cells (Lin^-^ CD122^+^ NK1.1^+^ DX5^+^) in the blood of *Casp9*^fl/fl^ and *Casp9*^fl/fl^ *Tie2-Cre*^+^ mice, in wildtype, *Ripk3*^*-/-*^ or *Casp1/11*^*-/-*^ background. Error bars indicate mean +/- S.D. of 3-10 mice, representative of at least two independent experiments. P-values calculated with two-way ANOVA followed by Tukey post hoc tests.

NK cells develop in the bone marrow from lymphoid progenitors, and commit to the NK cell lineage upon expression of CD122, a subunit of the receptor for interleukin-15 [36, 37]. These NK progenitors then acquire the expression of the NK cell lineage markers NK1.1 and DX5. The final stages of NK cell maturation are characterized by the ordered acquisition of CD11b and CD43 expression, and loss of CD27 expression [36]. This maturation process correlates with acquisition of NK cell effector functions [36]. In caspase-deficient mice, although commitment to the NK cell lineage in the bone marrow appeared to be intact, we observed reduced numbers of maturing NK cells as early as the CD11b^-^ CD43^-^ stage, in both the bone marrow and the spleen (**Figure 1D, E**). Frequencies of the different subsets suggest a relative blockade at the transition between the CD43^-^ and CD43^+^ stages, and this is more pronounced in caspase-3/7 deficient mice than in caspase-9 knockout (**Figure 1D, F**). A similar blockade is observed when maturation is monitored based on the induction of CD11b and loss of CD27 expression (**Figure 1D, Supplementary Figure 1C**). The expression of other cell surface markers, including activating and inhibitory receptors, is comparable in caspase sufficient and deficient mice (**Figure 1G**).

Besides caspase-dependent apoptosis, multiple mechanisms of regulated cell death exist. The functional crosstalk between these pathways determines by which molecular mechanism a cell dies [38]. Knocking out a central regulator of apoptosis may relieve inhibition on another cell death mechanism, as shown for example by the induction of RIPK3-dependent necroptosis during embryonic development of mice lacking caspase-8 [39, 40]. To test whether caspase-9 deficiency could result in the activation of RIPK3-mediated necroptosis or Caspase-1/11-mediated pyroptosis, and whether such alternative cell death could account for NK cell defects, we crossed *Casp9*^*fl/fl*^ *Tie2-Cre*^*+*^ mice to *Ripk3*^*-/-*^ [41] or *Casp1/Casp11*^*-/-*^ mice [42], respectively. In both cases, the absence of central effectors of necroptosis or pyroptosis failed to restore NK cell percentages in mice lacking caspase 9 (**Figure 1H**).

### NK cells are functionally defective in caspase-deficient mice

NK cell functional properties include cytotoxicity and cytokine production [32]. To evaluate NK-cell specific cytotoxicity *in vivo*, we differentially labeled splenocytes from wildtype and *B2m*^*-/-*^ mice [43], mixed them in a ∼50:50 ratio, and injected them in caspase-sufficient or deficient mice. Twelve hours later, we measured the ratio of persisting cells in the spleen (**Figure 2A**). Because *B2m*^*-/-*^ cells lack the β-2-microglobulin subunit of MHC class I, they are susceptible to NK cell-mediated cytotoxicity *in vivo* [44], and only ∼10% of them survive in caspase-sufficient mice. In contrast, ∼40% *B2m*^*-/-*^ cells persist in *Casp9*^*fl/fl*^ *Tie2-Cre*^*+*^ and in *Casp3*^*fl/fl*^ *Casp7*^*-/-*^ *Tie2-Cre*^*+*^ mice (**Figure 2A-B**), demonstrating altered NK cell cytotoxicity *in vivo*.

**Figure 2.**
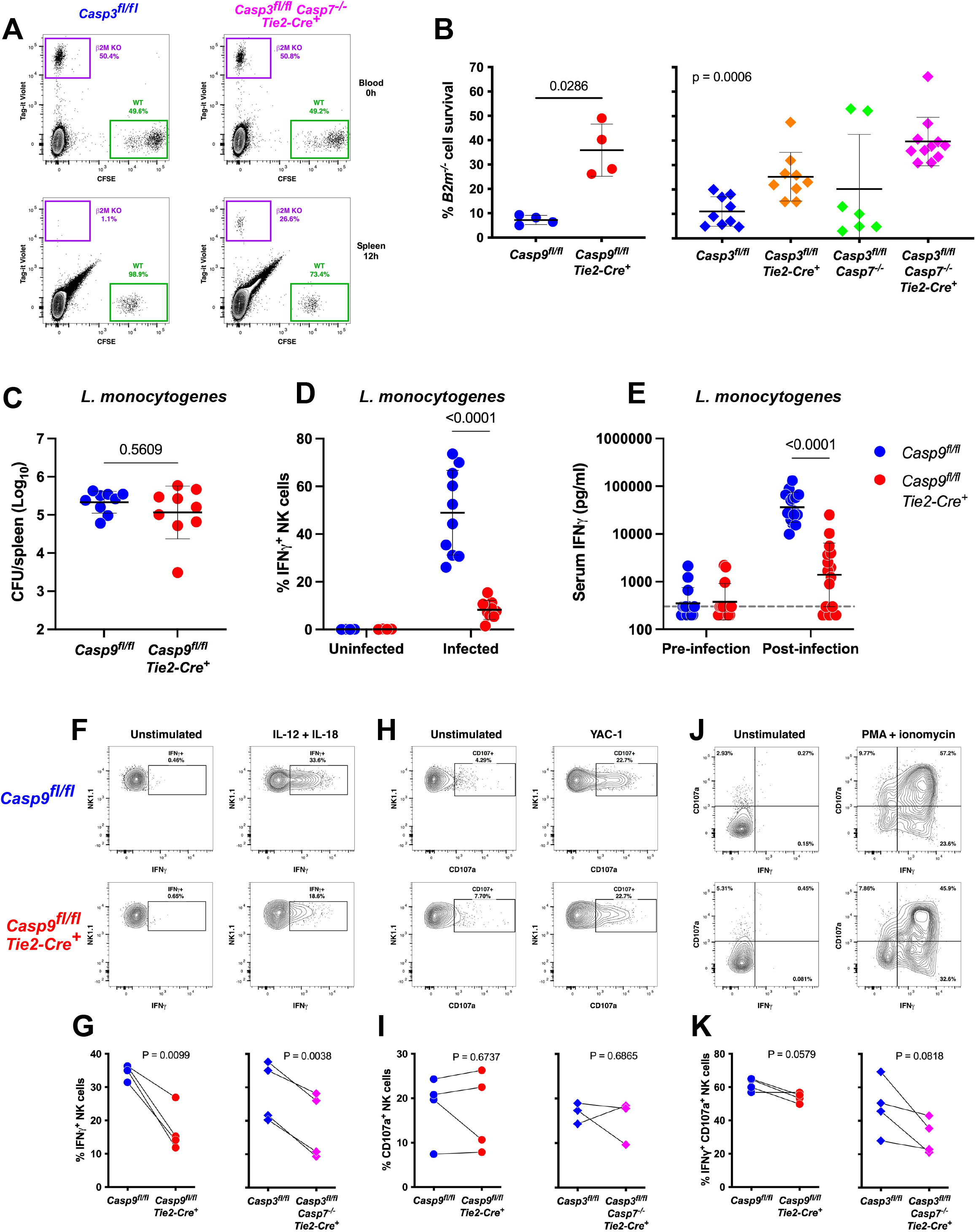
Defective NK cell function in caspase-deficient mice, *in vivo* and *in vitro*. (**A**) Representative visualization by flow cytometry of *in vivo* NK cell mediated cytotoxicity. Violet-labeled *B2m*^*-/-*^ splenocytes (targets for NK cell lysis) and CFSE-labeled wildtype splenocyte (non-target controls) were injected in a 1:1 ratio in recipient mice. The violet:CFSE ratio was determined in the blood right after injection and in the spleen 12 hours later. (**B**) Percentage of *B2m*^*-/-*^cells surviving NK cell cytotoxicity in vivo, calculated based on the violet:CFSE ratio at 12 hours, in the spleen of *Casp9*^fl/fl^ *Tie2-Cre*^+^, *Casp3*^fl/fl^ *Casp7*^-/-^ *Tie2-Cre*^+^ and respective control mice. Error bars indicate mean +/- S.D of N=4-11 mice/group. P-values calculated with Mann-Whitney or Kruskal-Wallis tests. (**C**-**E**) *Casp9*^fl/fl^ *Tie2-Cre*^+^ and control mice were infected intravenously with *L. monocytogenes* (∼10^4^ CFU/mouse) and analyzed 24 hours later. (**C**) Bacterial burden in the spleen measured by colony forming unit assay (N=9). (**D**) Frequency of IFNγ-producing NK cells (TCRβ^-^ NK1.1^+^), measured by intracellular cell staining after four hours of *ex vivo* culture with Brefeldin A and Monensin (N=4-10). (**E**) Serum concentration of IFNγ measured by ELISA before and after infection (N=13/15). The dashed line indicates limit of detection. P-values calculated with Mann-Whitney (**C**) or two-way ANOVA followed by Tukey post hoc tests (**D, E**). (**F-K**) Splenocytes from *Casp9*^fl/fl^ *Tie2-Cre*^+^, *Casp3*^fl/fl^ *Casp7*^-/-^ *Tie2-Cre*^+^ and respective control mice were stimulated with IL-12 and IL-18 (**F, G**), by YAC-1 cells (**H, I**) or with PMA and ionomycin (**J, K**), and activation was determined by flow cytometry based on IFNγ expression and CD107a exposure. (**F, H, J**) Representative flow plots. (**G, I, K**) Quantification of four independent experiments. P-values calculated with paired t tests.

On the other hand, NK cells are the main cellular source of IFNγ in response to *in vivo* infection with *Listeria monocytogenes* [45]. Following infection of *Casp9*^*fl/fl*^ and *Casp9*^*fl/fl*^ *Tie2-Cre*^*+*^ mice with *L. monocytogenes*, the bacterial burden was comparable between groups after 24 hours (**Figure 2C**). However, the frequency of IFNγ-producing NK cells (**Figure 2D**) as well as the serum concentration of IFNγ (**Figure 2E**) were significantly reduced in infected *Casp9*^*fl/fl*^ *Tie2-Cre*^*+*^ mice compared to their littermate controls.

*In vitro*, we stimulated splenocytes with the IL-12/IL-18 cytokine cocktail to induce IFNγ production, with the MHC class I deficient YAC-1 cells to induce NK cell degranulation, or with phorbol 12-myristate 13-acetate (PMA) and ionomycin as a strong activator. We used flow cytometry to measure IFNγ production and CD107a exposure (a marker of degranulation indicative of cytotoxic activity). We found that a lower frequency of NK cells produced IFNγ following IL-12/IL-18 treatment in the absence of caspases (**Figure 2F-G**). However, the response to YAC-1 cells and PMA/ionomycin was unaffected in NK cells from caspase-deficient mice (**Figure 2H-K**). Together, these results suggest that the reduced *in vivo* cytotoxicity is likely due to the lower number of NK cells (**Figure 1B-C**), while the defective IFNγ response can be attributed to reductions in NK cell numbers, as well as in lower responsiveness on a per-cell basis.

### NK cell deficiency is dependent on cGAS/STING, but independent of type I IFNs

Caspase-9 or caspases-3/7 deficiency results in constitutive activation of the type I IFN response and expression of interferon-stimulated genes (ISGs), resulting from mtDNA-mediated cGAS/STING activation [20, 21]. To test whether constitutive activation of the cGAS/STING pathway contributes to NK cell deficiency, we crossed *Casp9*^*fl/fl*^ *Tie2-Cre*^*+*^ mice with *Cgas*^*-/-*^ [46] or *Sting1*^*gt/gt*^ mice [47]. cGAS- or STING-deficiency both restored the baseline level of expression of *Isg15* in the absence of caspase-9 (**Figure 3A**). They also resulted in a partial, but significant, rescue of NK cell frequencies in the blood (**Supplementary Figure 2A**) and spleen of *Casp9*^*fl/fl*^ *Tie2-Cre*^*+*^ mice (**Figure 3B**). When comparing absolute cell counts in the spleen, cGAS or STING-deficiency entirely rescued NK cell numbers (**Supplementary Figure 2B**), and this could be attributed to a slight overall increase in spleen cellularity in double knockout mice (**Supplementary Figure 2C**). We also evaluated the functional properties of NK cells *in vivo* in *Casp9*^*fl/fl*^ *Cgas*^*-/-*^ *Tie2-Cre*^*+*^ mice. Upon *L. monocytogenes* infection, NK cells remained functionally defective in the double knockout mice as shown by their reduced IFNγ production (**Figure 3C**).

**Figure 3.**
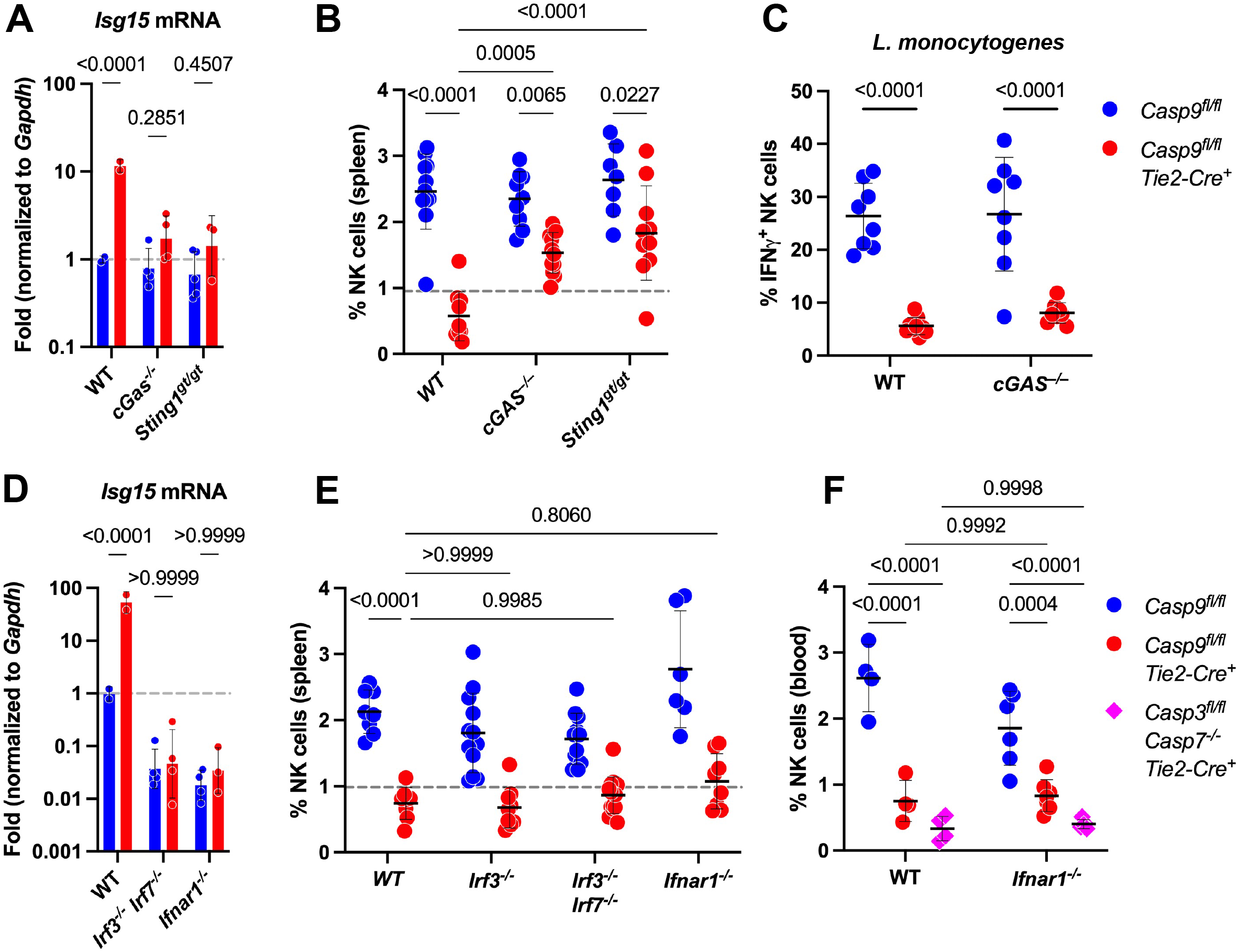
NK cell defect is partially dependent on cGAS/STING, but independent of type I IFN signaling. (**A, B**) Expression level of the mRNA encoding *Isg15* (**A**) and frequency of NK cells (Lin^-^ CD122^+^ NK1.1^+^ DX5^+^) in the spleen (**B**) of *Casp9*^fl/fl^ and *Casp9*^fl/fl^ *Tie2-Cre*^+^ mice, in wildtype, *cGAS*^*-/-*^ and *Sting1*^*gt/gt*^ background (N=7-12 mice/group, including males and females). *Isg15* was measured by realtime RT-PCR in white blood cells (N=2-3 mice/group), and normalized to *Gapdh*. The dashed line indicates 1 (**A**) or the mean + S.D. of *Casp9*^fl/fl^ *Tie2-Cre*^+^ mice (**B**). (**C**) Frequency of IFN γ -producing NK cells in the spleen of *Casp9*^fl/fl^ and *Casp9*^fl/fl^ *Tie2-Cre*^+^ mice, in wildtype or *cGAS*^*-/-*^ background, 24 hours after infection with *L. monocytogenes* (N=8-9 mice per group). (**D, E**) Expression level of the mRNA encoding *Isg15* (**D**) and frequency of NK cells in the spleen (**E**) of *Casp9*^fl/fl^ and *Casp9*^fl/fl^ *Tie2-Cre*^+^ mice, in wildtype, *Irf3*^*-/-*^, *Irf3*^*-/-*^*Irf7*^*-/-*^ and *Ifnar1*^*-/-*^ background (N=6-14 mice/group). (**F**) Frequency of NK cells in the blood of *Casp9*^fl/fl^, *Casp9*^fl/fl^ *Tie2-Cre*^+^ and *Casp3*^fl/fl^ *Casp7*^-/-^ *Tie2-Cre*^+^ mice in wildtype and *Ifnar1*^*-/-*^ background (N=4-7 mice/group). P-values calculated with two-way ANOVA followed by Šidák (**A, D**) or Tukey (**B, C, E, F**) post hoc tests.

We next generated caspase-9 deficient mice that also lack IRF3 and IRF7 – two transcription factors essential for type I IFN production [48, 49] – or the type I IFN receptor IFNAR1 [50]. In these mice, like in the absence of cGAS and STING, *Isg15* expression was restored to baseline levels in the absence of caspase-9 (**Figure 3D**). However, the NK cell frequencies remained defective in caspase-9 deficient mice lacking essential components of type I IFN signaling (**Figure 3E**). We also crossed mice lacking caspases 3 and 7 to the IFNAR1-deficient background, and the analysis of these mice confirmed that the NK cell defect is independent of type I IFNs (**Figure 3F**). Overall, these results provide *in vivo* genetic evidence that NK cell defects in caspase-deficient mice are not a direct consequence of constitutive expression of type I IFNs and ISGs. The reduction in NK cell frequency results in part from a non-canonical (i.e. type I IFN-independent) activity of the cGAS/STING pathway, while the defective production of IFNγ is independent of cGAS and STING.

### NK cell deficiency is cell-extrinsic

The *Tie2-Cre* transgene produces efficient deletion of floxed alleles in all immune cells [21, 35, 51] and in endothelial cells [35], as well as mosaic deletion in diverse tissues. The reduced NK cell frequency observed following transplantation of caspase-9 deficient hematopoietic progenitors in recipient mice (**Supplementary Figure 1E**) demonstrates the hematopoietic origin of the defect. To determine whether this origin is intrinsic to NK cells, we deleted caspase-encoding genes specifically in NK cells, using *Ncr1*^*iCre*^ knockin mice [52]. The resulting mice display efficient and selective deletion of the floxed *Casp9* or *Casp3* alleles in NK cells (**Figure 4A, C**). However, the abundance of NK cells was not affected by these NK cell-specific deletions of caspase-encoding genes (**Figure 4B, D**), suggesting a cell extrinsic origin to the phenotype.

**Figure 4.**
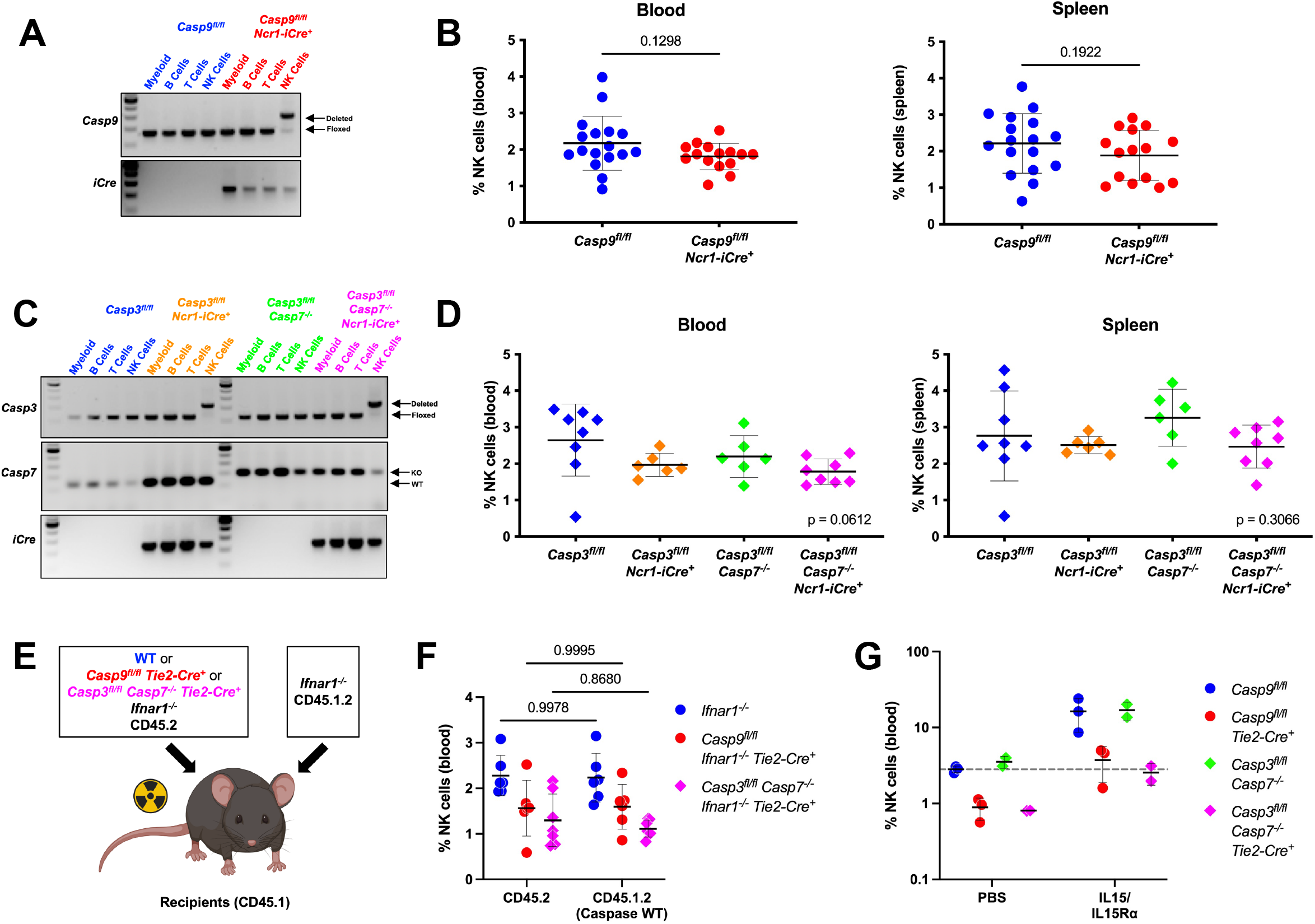
NK cell deficiency is a cell extrinsic phenotype of hematopoietic origin. (**A**-**D**) Genomic PCR visualizing the deletion of the floxed *Casp9* (**A**) or *Casp3* (**C**) alleles in immune cell populations isolated from the spleen of the indicated mice. Frequency of NK cells (Lin^-^ CD122^+^ NK1.1^+^ DX5^+^) in the blood of *Casp9*^fl/fl^ *Ncr1-iCre*^*+*^ (**B**), *Casp3*^fl/fl^ *Casp7*^-/-^ *Ncr1-iCre*^*+*^ (**D**) mice, and their respective controls. Error bars indicate mean +/- S.D of N=6-17 mice/group. P-values calculated with Mann-Whitney (**B**) or Kruskal-Wallis tests (**D**). (**E, F**) Schematic representation of the generation of mixed bone marrow chimeras (**E**). Frequency of NK cells (Lin^-^ NK1.1^+^) among CD45.2 cells with the indicated genetic modifications of caspase-encoding genes, and among wildtype CD45.1.2 cells from the corresponding mice (**F**). All transplanted cells are in *Ifnar1*^*-/-*^ background. N=6-7 mice per group, representative of three independent experiments. P-values calculated with two-way ANOVA followed by Šidák post hoc tests. (**G**) NK cell frequencies (Lin^-^ CD122^+^ NK1.1^+^ DX5^+^) in the blood of mice after treatment with PBS or recombinant IL-15/IL-15Rα treatment. Mean +/- S.D of N=2-3 mice/group. The dashed line indicates the mean of PBS-treated *Casp9*^*flfl*^ mice.

To further test this possibility, we generated mixed bone marrow chimeras, in which wildtype hematopoietic cells develop in the presence of caspase-deficient cells. Because caspase-deficiency, and the resulting type I IFN response, alters the fitness of hematopoietic stem cell function in mixed transplantation assays [20, 53], we used donor bone marrow cells from mice in *Ifnar1*^*-/-*^ genetic background (**Supplementary Figure 3**). Specifically, *Ifnar1*^*-/-*^ CD45.2, *Casp9*^*fl/fl*^ *Ifnar1*^*-/-*^ *Tie2-Cre*^*+*^ CD45.2, or *Casp3*^*fl/fl*^ *Casp7*^*-/-*^ *Ifnar1*^*-/-*^ *Tie2-Cre*^*+*^ CD45.2 bone marrow cells were mixed with *Ifnar1*^*-/-*^ CD45.1.2, and transplanted into irradiated wildtype CD45.1 recipient mice (**Figure 4E**). We observed that caspase-sufficient CD45.1.2 cells, when developing in the presence of caspase-deficient CD45.2 cells, acquire NK cell deficiency comparable to the caspase-deficient cells themselves (**Figure 4F**). These results demonstrate the cell extrinsic nature of the NK cell defect in caspase-deficient mice. They further show that caspase deficient hematopoietic cells, other than NK cells, harbor an activity that interferes with wildtype NK cell development in a dominant manner.

IL-15 is the essential cytokine supporting NK cell differentiation and homeostasis, and we wanted to test whether exogenous IL-15 could rescue NK cell development in caspase-deficient mice. Therefore, we treated *Casp9*^*fl/fl*^ *Tie2-Cre*^*+*^, *Casp3*^*fl/fl*^ *Casp7*^*-/-*^ *Tie2-Cre*^*+*^, and their respective littermates with recombinant IL-15/IL-15Rα complex. This treatment resulted in the expansion of NK cells by a similar ∼3-5 fold in all groups, bringing NK cell numbers in knockout mice to levels comparable to their untreated wildtype littermates mice (**Figure 4G**). This result shows that caspase-deficient NK cells can reach normal levels under adequate experimental conditions, and is compatible with the cell-extrinsic nature of their defects.

## Discussion

Genetic deficiencies in effectors of apoptotic cell death generally produce autoimmune or inflammatory phenotypes. Causes include dysregulated immune cell homeostasis leading to hyperplasia [5, 6], failure of immune response resolution resulting in prolonged immune cell activation [54], or induction of an alternative cell death mechanism with pro-inflammatory characteristics [38]. In this context, our observation of NK cell defects in pro-apoptotic caspase-deficient mice is unusual, since it results in a decrease in NK cell number and reduced responsiveness to stimulation, rather than the expected accumulation of “non-dying” cells.

Previous research showed that Bax/Bak deficiency, but not caspase-9 deficiency, resulted in immune cell hyperplasia [11]. This observation suggested the existence of a caspase-9-independent cell death mechanism downstream of Bax/Bak; this is confirmed by our results using conditional *Casp9* knockout mice. Furthermore, we show that this mechanism is also independent of the downstream effectors, caspases-3 and -7. The precise mechanisms of caspase-independent cell death downstream of Bax/Bak remain unknown, but recent reports have shown that RIPK3-dependent necroptosis is activated in the absence of caspase-9, under specific experimental conditions [55, 56]. However, our genetic *in vivo* data rules out alternative death by RIPK3-mediated necroptosis or caspase-1/11-dependent pyroptosis, which may indirectly affect NK cells.

Caspase-9 is activated following outer membrane mitochondrial permeabilization (MOMP) in cells initiating apoptotic cell death [57]. It is also activated in cells undergoing mild stress, resulting in permeabilization of a minority of mitochondria, a process known as miMOMP. Although caspase-9 activation is not lethal in cells undergoing miMOMP, it nevertheless has functional consequences on cell physiology [58, 59]. miMOMP in the absence of active caspase-9 results in the leakage of mtDNA through Bax/Bak [22, 23], activation of the pro-inflammatory cGAS/STING pathway and type I IFN production [20, 21]. Consequently, the type I IFN response is constitutively active in caspase-9 deficient mice, but this IFN response is not responsible for NK cell deficiency. Indeed, knocking out the IRF3/IRF7 transcription factors, or the IFNAR1 receptor, doesn’t rescue NK cell numbers in mice lacking caspase-9. However, cGAS or STING deficiency partially rescues NK cell frequencies, linking the phenotype to DNA sensing and, likely, to mitochondrial permeability and mtDNA release. Together, our observations demonstrate that in cells that undergo MOMP but fail to activate apoptosis, a type I IFN-independent STING activity alters NK cells in a cell-extrinsic manner. Besides type I IFNs, STING triggers NF-κ B signaling and ATG5-dependent autophagy [60, 61], and both mechanisms could impact NK cells.

Besides experimental conditions in which caspase activity is blocked, the mtDNA/cGAS/STING axis is activated in a number of pathological conditions [62]. These include infections, such as with dengue virus or particular strains of *Mycobacterium tuberculosis*, in which type I IFN expression is triggered by infection-induced mitochondrial stress and mtDNA, rather than by direct pathogen sensing [25, 26, 28]. Mitochondrial stress and mtDNA leakage also activate cGAS/STING in inflammatory and neurodegenerative diseases, such as systemic lupus erythematosus and amyotrophic lateral sclerosis [30, 63]. In these conditions, studies of the mtDNA/cGAS/STING axis have focused primarily on the production of type I IFN and other inflammatory mediators – a potential impact on NK cell development, homeostasis and function, as well as the consequences on disease outcomes, will need to be considered in future research.

## Acknowledgments

We thank Drs. Andrew Oberst, Daniel Stetson, Tadatsugu Taniguchi, Eric Vivier and Irving Weissman for sharing mice; Dr. Deborah Banker for manuscript editing; and Silvia Christian for administrative support. This work was supported by the Bezos family, the Lupus Research Alliance and the National Institute of Allergy and Infectious Diseases of the National Institutes of Health (R21 AI138011). This research was also supported by Shared Resources (Comparative Medicine, Flow Cytometry) of the Fred Hutchinson Cancer Research Center/University of Washington Cancer Consortium (P30 CA015704).

## Material and Methods

### Mice

Mice with Tie2-Cre (E+H) [35] -mediated conditional deletion of caspase-9, or of caspase-3 in caspase-7 knockout background [21, 64]; transgenic H2K-*Bcl2* mice [4]; caspase-1/11 [42], RIPK3 [41], β_2_ micro-globulin [43], cGAS [46], STING [47], IRF3 [48], IRF7 [49], and IFNAR1 [50] -deficient mice; and *Ncr1*-iCre knockin mice [52] have been previously reported. *Casp9*Δ/ Δ were generated by Cre-mediated germline deletion of the floxed allele of *Casp9*^*fl/fl*^ mice, taking advantage of mosaic deletion in a *Tie2-Cre*^*+*^ female, and the *Cre* transgene was bred out. All experiments were performed with littermate controls and included males and females. Experiments were performed in compliance with Fred Hutch’s Institutional Care and Use Committee protocol 50941.

### Fetal liver chimeras and mixed bone marrow chimeras

Recipient CD45.1 female mice (obtained from Charles Rivers) were irradiated in a Cs irradiator (800 cGy gamma rays in a Cesium-137 irradiator). Four hours later, cells were injected by retroorbital intravenous route under Isoflurane anesthesia. Mice were provided with antibiotic in drinking water (dose) for 2 weeks after transplantation.

For fetal liver chimeras, fetal tissues were harvested from embryos at E16.5-E18.5 and genotyped, single cell suspension of the liver were prepared, and *Casp9*^*+/+*^ and *Casp9*^Δ/ Δ^ cells from littermate were injected (one liver for 10 recipient mice). For mixed bone marrow chimeras, bone marrow cells were harvested from donors in CD45.2 and CD45.1.2 background, mixed in a ∼1:1 ratio and a total of 2 million cells were injected intravenously.

### Flow cytometry

Blood was prepared by red blood cell lysis in Ammonium-Chloride-Potassium (ACK) solution. Single cell suspensions were prepared from the spleen or bone marrow, treated by ACK lysis, and resuspended in FACS buffer (phosphate buffered saline supplemented with 1% bovine serum albumin, 2 mM EDTA and 0.02% NaN_3_). All staining were performed in FACS buffer and dead cells were counterstained with 7-amino-actinomycin D (7-AAD, Biolegend). Before analysis, cells were pre-gated on live (7-AAD^-^) CD45^+^ singlet cells. The following antibodies were used (all from Biolegend, unless otherwise specified): CD4-PE (clone GK1.5), CD8α-AF700 (53-6.7), CD11b-AF700 and -BV785 (M1/70), CD19-APC and -BV510 (6D5), CD27-PacificBlue (LG.3A10), CD43-FITC (S7, BD Pharmingen), CD45-APC-Fire750 (30-F11), CD45.1-PE (A20), CD45.2-APC (104), CD49b-PE (DX5), CD107a-APC (1D4B), CD122-biot (TM-b1), CD132-PE (TUGm2), CD192/CCR2-biot (SA203G11), CD314/NKG2D-FITC (C7), CD335/NKp46-PE (29A1.4), IFNγ-FITC (XMG1.2), Ly-49A-FITC (Ye1/48.10.6), Ly-49D-PE (4E5), Ly-49G2-FITC (eBio4D11, eBioscience), Ly-6C-FITC (HK1.4), Ly-6G-PacificBlue (1A8), NK1.1-PE-Cy7 (PK136), NKG2A-PE (16a11, eBioscience), TCRβ-APC and -PacificBlue (H57-597), TER-119-APC. Streptavidin-BV605 (Biolegend) was used as a secondary reagent with biotinylated antibodies.

In most experiments, the lineage cocktail contained Ter119, TCRβ and CD19-APC antibodies, and NK cells were identified as 7AAD^-^ CD45^+^ Lin^-^ CD122^+^ NK1.1^+^ DX5^+^ cells.

Results were analyzed with Flow Jo (version 10.7.1, Becton Dickinson & Company).

### *In vivo* NK cell cytotoxicity assay

Recipient mice were pre-treated with poly-(I:C) (100 μg/mouse, from Sigma-Aldrich), injected intraperitoneally ∼16 hours in advance. Splenocytes from donor wildtype mice or from *B2m*^*-/-*^ mice were labeled with carboxyfluorescein diacetate succinimidyl ester (CFSE, Biolegend) or with Tag-It Violet (Biolegend), mixed in a 1:1 ratio, and injected intravenously (∼1 × 10^6^ cells/mouse) into recipient mice. Blood was collected immediately to confirm the input ratio. Mice were killed 12 h later and a single cell suspension of the spleen was prepared and analyzed by flow cytometry. At least 500 CFSE^+^ events were acquired for each sample. The proportions of violet *B2m*^-/-^ cells (targets of NK cells) and CFSE^+^ wildtype cells (internal controls) were measured and the percentage of survival to specific NK cell lysis was calculated as:

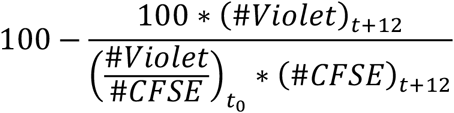

Where # indicates the number of events acquired in the Violet^+^ and CFSE^+^ gates, respectively.

### *In vivo* Listeria infection

Mice were infected with ∼10^4^ colony-forming units (CFU) of L. monocytogenes (strain 10403S) by intravenous injection, and serum and spleens were collected 24 hours later. Splenocytes from uninfected or infected mice were incubated at 37C° in 5% CO2 for 4 h in medium containing Monensin and Brefeldin A solutions (Biolegend). Cells were then stained for surface antigens (TCRβ-PacificBlue and NK1.1-PE), permeabilized using Biolegend’s Fixation and Perm Wash buffers, and stained intracellularly with an anti-IFNγ-FITC antibody. Serum concentrations of IFNγ were measured using ELISA MAX Standard set (Biolegend), following the manufacturer’s instructions.

### *In vitro* NK cell stimulation

Splenocytes were incubated at 37°C in 5% CO2 for 4 h in medium containing anti-CD107a-APC antibody (1 μg/ml), Monensin and Brefeldin A solutions (Biolegend), under one of the following stimulation conditions: (1) untreated, (2) treated with recombinant mouse IL-12 (5 ng/ml) and IL-18 (25 ng/ml, both from Peprotech), (3) mixed with YAC-1 cells in a 0.4:1 (YAC-1:splenocyte) ratio, or (4) treated with phorbol 12-myristate 13-acetate (PMA, 2.5 μg/ml) and ionomycin (0.25 ng/ml, both from Sigma-Aldrich). After incubation, cells were stained intracellularly for IFNγ expression, as described above, and analyzed by flow cytometry.

### IL-15 treatment

Mice were treated with IL-15/IL-15Rα complex as previously reported [65]. Briefly, recombinant mouse IL-15Rα-Fc and recombinant human IL-15 were purchased from R&D Systems. For each mouse, a mix 2.5 μg of IL-15 and 15 μg of IL-15Rα-Fc was resuspended in 200 μl PBS and incubated for 30 minutes at 37°C. The complex was injected intraperitoneally twice (two days apart), and analysis was performed three days after the second injection.

### Gene Expression Analysis

RNA was extracted (RNeasy Plus mini kit, QIAGEN) from white blood cells, reverse transcribed (SuperScript III, Invitrogen) and the expression of specific genes was analyzed by SybrGreen-based real time PCR (iTaq Universal SYBR Green Supermix, BIO-RAD) on a CFX Connect Real-Time System (BIO-RAD).

The following primers were used:

*Isg15* forward: CAAGCAGCCAGAAGCAGACT

*Isg15* reverse: CCCAGCATCTTCACCTTTAGG

*Gapdh* forward: GGTGTCTTCACCACCATGGA

*Gapdh* reverse: CGGAGATGATGACCCTTTTG

### Statistical analyses

All statistical analyses were performed with GraphPad Prism (version 9). Means of experimental groups were compared using the Mann-Whitney or paired t tests (two groups), or the Kruskal-Wallis test (four groups) test. When two parameters are involved, the two-way ANOVA test was used, followed by Tukey or Šidák post hoc tests for multiple comparisons.

---

**Supplementary Figure 1.**
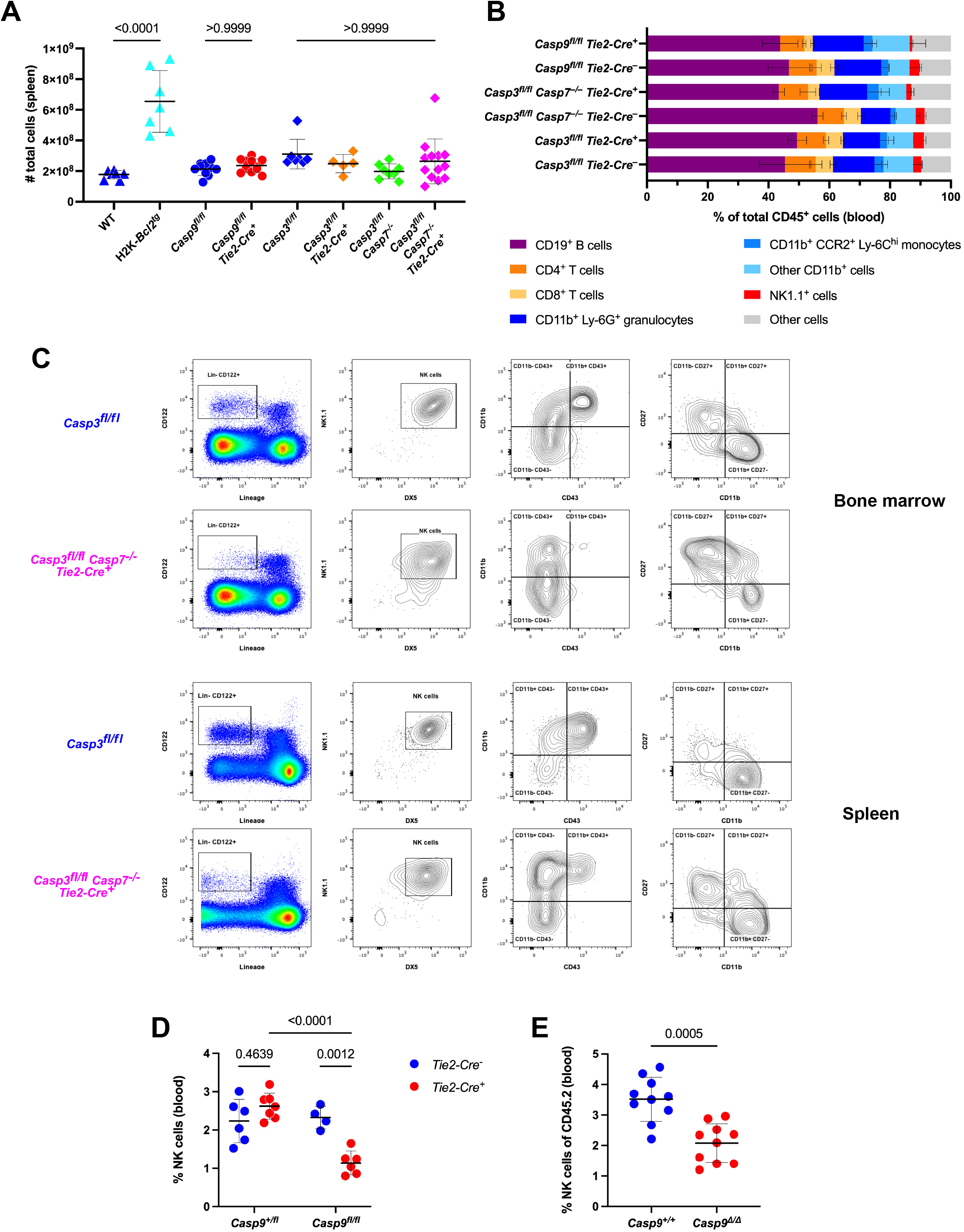
(**A**) Absolute cell numbers, after red blood cell lysis, in the spleen of H2K-*Bcl2* transgenic, caspase deficient and respective littermate control mice. Error bars indicate mean +/- S.D. of N=5-13 mice per group. P-values calculated with the Kruskal-Wallis test. (**B**) Frequencies of immune cell lineages in the blood of caspase deficient and littermate control mice. Error bars indicate mean +/- S.D. of N=3-4 mice/group. (**C**) Representative flow plots illustrating the gating strategy used to identify NK cells, as well as the expression of maturation markers, in the bone marrow and spleen of *Casp3*^fl/fl^ and *Casp3*^fl/fl^ *Casp7*^-/-^ *Tie2-Cre*^+^ mice. (**D**) Frequencies of NK cells (Lin^-^ CD122^+^ NK1.1^+^ DX5^+^) in the blood of *Casp9*^+/fl^, *Casp9*^+/fl^ *Tie2-Cre*^+^, *Casp9*^fl/fl^ and *Casp9*^fl/fl^ *Tie2-Cre*^+^ littermates. Error bars indicate mean +/- S.D. of N=4-7 mice per group. P-values calculated with two-way ANOVA followed by Šidák post hoc tests. (**E**) Frequencies of NK cells (Lin^-^ CD122^+^ NK1.1^+^ DX5^+^) among CD45.2^+^ cells in the blood of CD45.1^+^ mice transplanted with fetal liver cells from *Casp9*^*Δ/Δ*^ or littermate *Casp9*^*+/+*^ embryos. Error bars indicate mean +/- S.D. of N=10 mice/group. P-value calculated with the Mann-Whitney test.

**Supplementary Figure 2.**
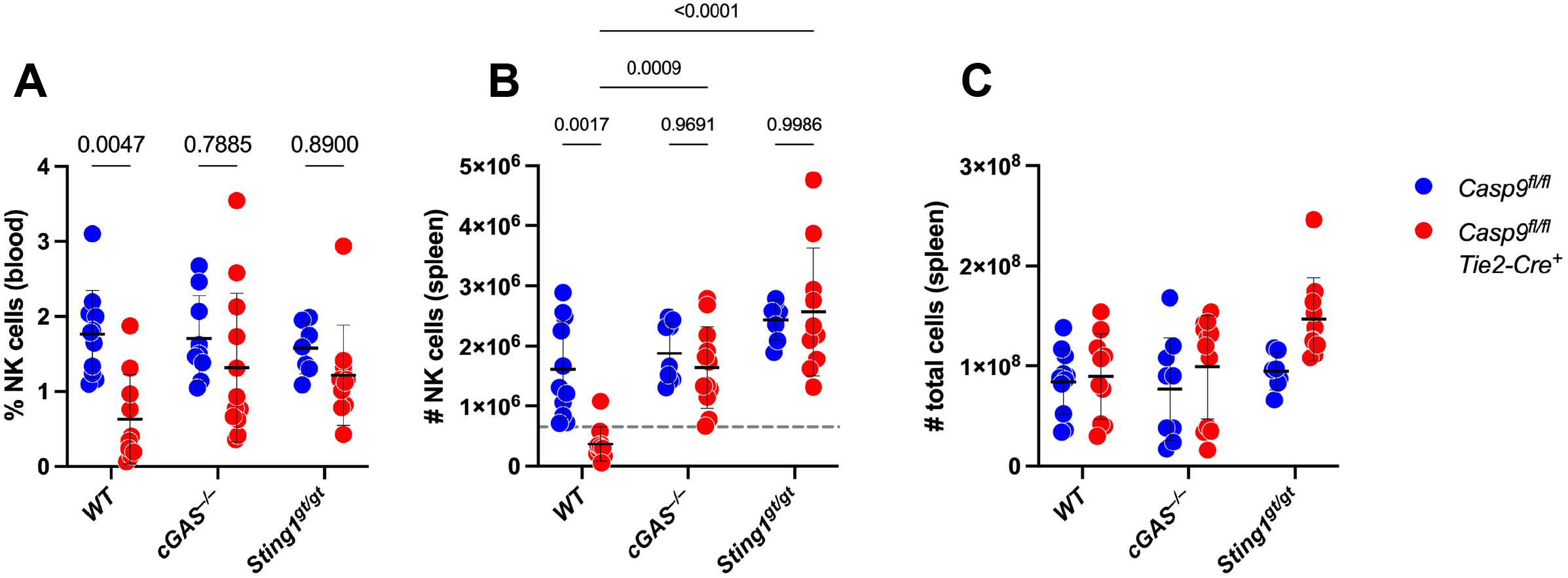
(**A, B**) Frequency in the blood (**A**) and absolute numbers in the spleen (**B**) of NK cells (Lin^-^ CD122^+^ NK1.1^+^ DX5^+^) in *Casp9*^fl/fl^ and *Casp9*^fl/fl^ *Tie2-Cre*^+^ mice, in wildtype, *cGAS*^*-/-*^ and *Sting1*^*gt/gt*^ background (N=7-12 mice/group). P-values calculated with two-way ANOVA followed by Tukey post hoc tests. (**C**) Total spleen cell numbers in the same spleens.

**Supplementary Figure 3.**
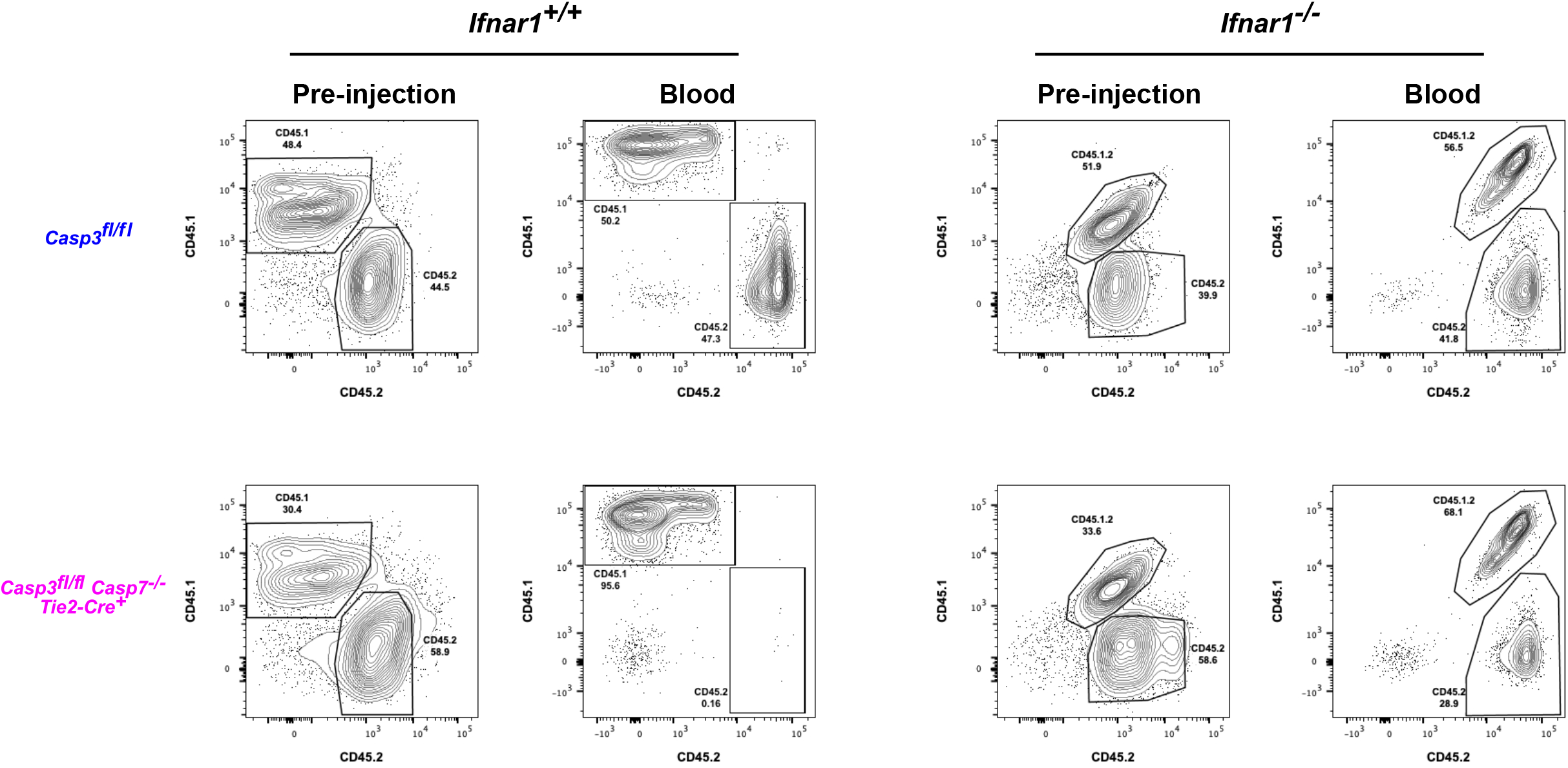
Flow plots illustrating the ratio of CD45.2 (caspase wildtype or knockout) and CD45.1 or CD45.1.2 (wildtype) prior to injection, or in the blood of recipient mice 6 weeks after injection. Mixed wildtype cells were CD45.1, while mixed *Ifnar1*^*-/-*^ cells were CD45.1.2.

